# An Optimized Circular Polymerase Extension Reaction-based Method for Functional Analysis of SARS-CoV-2

**DOI:** 10.1101/2022.11.26.518005

**Authors:** GuanQun Liu, Michaela U. Gack

**Affiliations:** Florida Research and Innovation Center, Cleveland Clinic, Port St. Lucie, Florida, USA

## Abstract

**SUMMARY:** Reverse genetics systems have been crucial for studying specific viral genes and their relevance in the virus lifecycle, and become important tools for the rational attenuation of viruses and thereby for vaccine design. Recent rapid progress has been made in the establishment of reverse genetics systems for functional analysis of SARS-CoV-2, a coronavirus that causes the ongoing COVID-19 pandemic that has resulted in detrimental public health and economic burden. Among the different reverse genetics approaches, CPER (circular polymerase extension reaction) has become one of the leading methodologies to generate recombinant SARS-CoV-2 infectious clones due to its accuracy, efficiency, and flexibility. Here, we report an optimized CPER methodology which, through the use of a modified linker plasmid and by performing DNA nick ligation and direct transfection of permissive cells, overcomes certain intrinsic limitations of the ‘traditional’ CPER approaches for SARS-CoV-2, allowing for efficient virus rescue. This optimized CPER system may facilitate research studies to assess the contribution of SARS-CoV-2 genes and individual motifs or residues to virus replication, pathogenesis and immune escape, and may also be adapted to other viruses.

## INTRODUCTION

Functional analysis of individual viral genes including embedded motifs and individual residues has been essential for understanding key functions of viruses such as viral entry, genome amplification, or escape from innate or adaptive immunity. Key to these studies has been the establishment of viral reverse genetics systems, which allow investigation of viral gene functions through mutagenesis [1, 2]. In addition, reverse genetics approaches for generating mutant recombinant viruses have become important for the rational design of replication-impaired, so-called “live-attenuated” viruses, which may represent vaccine candidates. Moreover, reverse genetics technologies enable studying viral evasion of antibody responses (*e.g*. by the coronaviral spike protein) and thereby aid in mRNA vaccine design [3]. Therefore, the development of efficient and accurate methodologies for generating viral infectious clones including recombinant mutant viruses has not only become an integral component of fundamental virology research, but also has great value for translational research and the design of novel vaccines [4].

SARS-CoV-2, a member of the large family of *Coronaviridae*, emerged in Wuhan, China, in late 2019 and then spread rapidly across the globe where it has caused substantial morbidity and mortality as well as severe economic losses [5]. SARS-CoV-2 is one of the largest RNA viruses. Its positive-sense genome is ~30 kb long and comprises a defined organization that encodes for ~30 gene products or proteins [6]. Since the emergence of SARS-CoV-2, rapid progress has been made in understanding how individual viral proteins or enzymes (*e.g*. spike protein or the RNA-dependent RNA polymerase) fulfill key functions in the viral lifecycle such as mediating virus entry and immune evasion or genome amplification. Studies to characterize viral proteins in isolation – either through ectopic expression in mammalian cells or by *in vitro* analysis following protein purification – have tremendously enhanced our understanding of how SARS-CoV-2 proteins function and provided important insight into their catalytic activities or interactions with host-cell factors or other viral proteins. However, the engineering of mutant recombinant viruses in which specific genes/residues are deleted or mutated has been essential for determining how relevant individual genes or specific motifs/residues are for virus infection, pathogenesis or immune evasion. The large genome size of SARS-CoV-2 has hampered the development of plasmid-based reverse genetics systems for this virus (and also other coronaviruses) that have been used for many other RNA viruses (*i.e*. influenza and flaviviruses) [4]. Therefore, bacterial artificial chromosome (BAC)-based technologies (typically used for mutagenesis of large DNA viruses such as herpesviruses), *in vitro* cDNA fragment ligation, and yeast-based synthetic biology approaches have been traditionally used for generating recombinant coronaviruses including SARS-CoV-2 [7–15].

In 2021, the adaptation of a circular polymerase extension reaction (CPER)-based approach, which has been successfully used for construction of flavivirus infectious clones [16], was reported for the generation of recombinant SARS-CoV-2 [17, 18]. Advantages of the CPER method include high-fidelity preservation of viral genome sequences with minimal or no unwanted mutations, as compared to the BAC and *in vitro* ligation methodologies which can introduce inexplicable insertions or deletions during bacterial propagation steps. Additionally, CPER allows for flexibility in viral sequence manipulation by PCR-based mutagenesis, while the BAC methodology relies on *de novo* assembly or homologous recombination in special bacterial systems. Furthermore, the straightforward and streamlined workflow of CPER allows for infectious clone construction in a single-tube reaction, which is in sharp contrast to BAC cloning and *in vitro* ligation of cDNA fragments that require cumbersome procedures and complex experimental techniques.

Integral to the CPER technology is PCR-based amplification of cDNA fragments that cover the complete viral genome (30 kb in the case of SARS-CoV-2) and carry overlapping sequences. With the use of a ‘linker’ fragment that connects the viral 5’ and 3’ untranslated regions (UTRs) with functional mammalian transcription initiation and termination elements, the individual cDNA fragments are extended in a single PCR reaction to assemble into a circularized full-length viral cDNA clone. The circularized cDNA clone is then delivered (typically by transfection) into mammalian cells, leading to the intracellular synthesis of viral genomic RNA and, ultimately, the production of infectious virus. Although the CPER platform has already greatly facilitated studies to functionally characterize SARS-CoV-2 genes and specific mutations, some intrinsic limitations still exist that hamper the robustness and efficiency of virus rescue.

Here, we report an optimized CPER methodology for reverse genetics engineering of SARS-CoV-2. Specifically, we utilized a modified linker plasmid, added a new step of ligating DNA nicks, and also applied direct transfection of the circularized infectious cDNA clone into highly permissive cells, which resulted in more rapid rescue of the virus and efficient viral yields.

## RESULTS

### Optimization of the CPER approach for efficient SARS-CoV-2 rescue

The CPER method builds principally on overlap extension PCR that fuses several double-stranded DNA (dsDNA) fragments containing 20-to 50-bp homologous ends into one large fragment [19]. Compared to the traditional overlap extension PCR, which uses a set of two distal primers to facilitate the generation of the combined fragment, CPER does not amplify fragments using such primers but instead utilizes an additional fragment that overlaps with the first and the last fragment to be joined, thereby circularizing the self-primed and extended dsDNA product. In CPER-based bacterial cloning, this additional fragment is typically a linearized plasmid vector generated by restriction digestion or PCR. As a result, the CPER product resembles a relaxed circular plasmid with staggered nicks which locate to the 5’ end of each strand of the individual fragment following the respective ‘round-the-horn’ amplification, as commonly seen in the QuikChange® approach of site-directed mutagenesis [20].

Adaptation of the CPER approach to *de novo* assembly of infectious clones for positive-strand RNA viruses is primarily achieved by substituting a linker fragment for the linearized vector used in CPER-mediated plasmid cloning. The design of the linker fragment draws inspiration from plasmid-launched mRNA synthesis driven by the mammalian RNA polymerase II (Pol II) promoter, as the genomes of several positive-sense RNA viruses including flaviviruses and coronaviruses contain a 5’ cap structure like cellular mRNAs and undergo cap-dependent translation. In addition to the Pol II promoter, the linker fragment also contains a polyadenylation signal for transcription termination and, importantly, a self-cleaving ribozyme sequence in front of the poly(A) signal to ensure accurate processing of the 3’ end of the RNA transcript to match the authentic viral genome sequence. Notably, while the linker fragment is usually cloned into a plasmid for long-term maintenance in *E. coli*, only the portion containing the mammalian transcription elements, but not the bacterial propagation cassettes, is amplified and used in CPER assembly.

Despite the successful adaptation of the CPER technology for the generation of infectious clones, CPER has a major intrinsic limitation, which is the presence of staggered nicks that impede efficient expression in mammalian cells. Whereas nicked plasmids are known to be seamlessly repaired upon transformation into *E. coli*, the precise fate of a circularized, nick-containing dsDNA inside a mammalian cell remains elusive. The presence of nicks in the template strand can cause Pol II pausing and likely also template misalignment, which may eventually lead to unwanted mutations [21]. In CPER-derived infectious clones, the circular template strand extended from each fragment contains a nick, which, depending on the genome segmentation scheme used for assembly, locates to different coding or noncoding regions of the viral genome. Although the sequence contexts in which the nicks situate may permit Pol II bypassing, how the template discontinuity affects the overall Pol II transcription efficiency, and hence the synthesis of full-length viral genomes, in mammalian cells remains unclear.

Another limitation of the current CPER approaches for SARS-CoV-2 rescue lies in the choice of cell lines for transfection of the CPER product [17, 18]. While HEK293-derived cell lines have been successfully used for reverse genetics systems for a variety of viruses from diverse families due to their robust transfectability, the use of HEK293 cells for SARS-CoV-2 rescue can be less efficient because of the unique cellular tropism of the virus and the critical host factors required for virus entry and replication. To date, three mammalian cell lines are commonly used for *in vitro* propagation of SARS-CoV-2 to high titers. These include Vero E6 (African green monkey kidney epithelial), Caco-2 (human colonic epithelial), and Calu-3 (human lung epithelial) cells. All three cell lines express the receptor for SARS-CoV-2, angiotensin-converting enzyme 2 (ACE2), while the latter two express also transmembrane serine protease 2 (TMPRSS2), a critical early entry cofactor [22]. In addition to priming direct cell membrane fusion, the presence of TMPRSS2 safeguards the integrity of the polybasic furin cleavage site in the viral spike gene, which is selectively deleted during serial passaging in Vero E6 cells due to viral host adaptation [23]. To this end, Vero E6 cells stably expressing human TMPRSS2 (Vero E6-TMPRSS2) have been widely used for the propagation of ancestral and emerging SARS-CoV-2 strains including the variants of concern (VOCs). More importantly, given the nature that Vero cells lack interferon (IFN) production [24], it remains the first-line cell system for generating and propagating recombinant mutant viruses that are attenuated through selective ablation of viral gene functions that evade or antagonize IFN-mediated antiviral innate immunity *(e.g*. SARS-CoV-2 papain-like protease (PLpro) which is an IFN antagonist [25, 26]).

Taking these limitations into account, we rationally optimized CPER for SARS-CoV-2 by adding new steps to seal the nicks in the CPER product and by using a modified linker plasmid as well as a different cell line for transfection of the CPER product (**Figure 1A**). Specifically, under the same genome segmentation scheme reported by Torii *et al*. [18], gel-purified viral cDNA fragments were phosphorylated at the 5’ end by using a T4 polynucleotide kinase. Equal molar amounts of the phosphorylated fragments were then subjected to CPER assembly using the cycling ‘condition 3’ as described previously [18]. Immediately before transfection, the nicks in the CPER product were sealed by using a high-fidelity and thermostable Taq DNA ligase that joins the extended 3’–OH terminus with its originating 5’–phosphorylated terminus, giving rise to a closed circular cDNA infectious clone. Then, the sealed CPER product was directly transfected into a monolayer of Vero E6-TMPRSS2 cells by using the *TransIT-X2* dynamic delivery system (**Figure 1A**). Furthermore, to ensure efficient Pol II termination and to prevent Pol II read-through in the linker region, which may confound ribozyme processing at the transcript 3’ end or interfere with new transcription initiation, we also replaced the ‘spacer’ sequence that is located between the poly(A) signal and CMV enhancer/promoter with a functional Pol II transcriptional pause signal from the human α2 globin gene known to minimize promoter crosstalk [27]. The resultant linker sequence was assembled with ampicillin resistance and origin of replication cassettes into a high-copy plasmid, named “pGL-CPERlinker” (**Figure 1B**).

**FIGURE 1.**
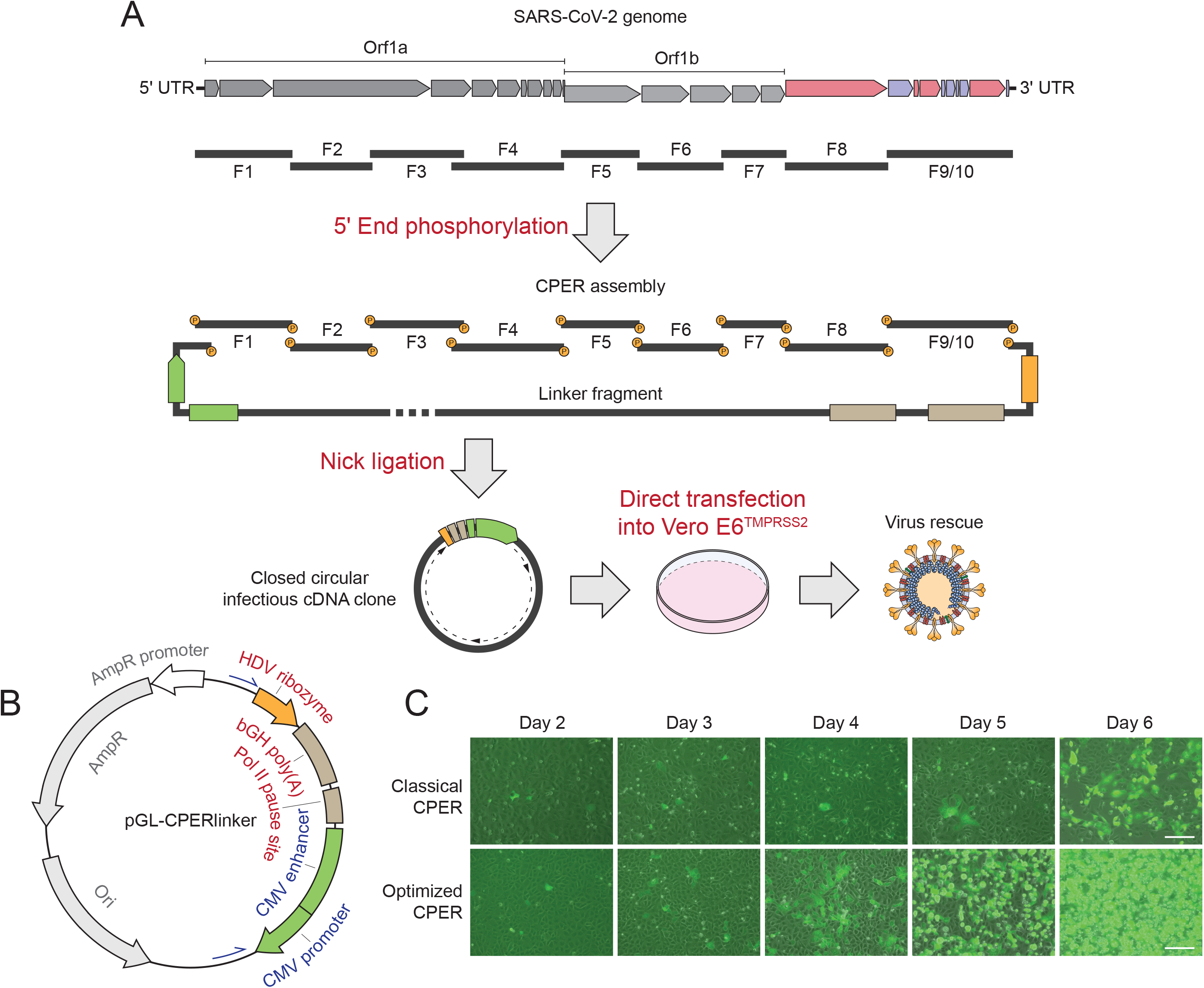
Generation of recombinant SARS-CoV-2 by using an optimized CPER methodology. (**A**) Schematic of the optimized CPER system that includes new or modified steps including 5’ end phosphorylation, nick ligation, as well as direct transfection of permissive cells with the CPER product. Specifically, the nine overlapping cDNA fragments (F1-F9/10) covering the full-length SARS-CoV-2 genome were phosphorylated at the 5’ end using T4 polynucleotide kinase (PNK) before being subjected to CPER assembly using a modified linker fragment (as illustrated in B). The circularized CPER product was then sealed at the staggered nicks by DNA ligation using HiFi *Taq* DNA ligase, and the closed circular infectious cDNA clone was transfected into Vero E6-TMPRSS2 cells for virus rescue. ‘P’ indicates phosphorylation. (**B**) Map of the linker plasmid (pGL-CPERlinker) in which the hepatitis delta virus (HDV) ribozyme, bovine growth hormone polyadenylation signal (bGH polyA), RNA polymerase II (Pol II) transcription pause signal, and human cytomegalovirus (CMV) enhancer and promoter were assembled together with the ampicillin resistance (AmpR) cassette and the origin of replication (Ori) derived from the pUC19 plasmid (NEB). (**C**) Comparison of the optimized CPER system with the original method as described by Amarilla *et al*. [17] by rescuing a GFP reporter virus. GFP-positive syncytia were evident as early as day 3 and day 5 post-transfection of the CPER product into Vero E6-TMPRSS2 cells, respectively. Scale bar, 100 μm.

Using the newly optimized CPER workflow, infectious virus generated using as a template a BAC construct encoding a GFP reporter SARS-CoV-2 [11] could be rescued as early as day 3 post-transfection, as evidenced by the formation of GFP-positive syncytia (**Figure 1C**). By day 5 post-transfection, massive cytopathic effects (CPE) could be observed. In comparison, successful virus rescue using the ‘classical’ CPER approach was not observed until day 5 post-transfection (**Figure 1C**), similar to previous reports [17, 18]. Therefore, the optimized CPER workflow can accelerate SARS-CoV-2 rescue by at least 2 days.

### Cloning-free SARS-CoV-2 rescue and characterization of the CPER-derived recombinant viruses

We also applied the optimized CPER approach to rescue SARS-CoV-2 from purified viral genomic RNA [17]. Adopting again the 10-fragment scheme reported by Torii *et al*. [18], we successfully achieved specific amplification of all fragments from the first-strand cDNA that was synthesized from purified viral genomic RNAs of three different virus strains, including the ancestral strain WA1 and two VOCs (*i.e*. Beta and Omicron) (**Figure 2A**). We also performed site-directed mutagenesis directly in the purified fragment #2 by overlap extension PCR using the pair of primers for fragment #2 amplification (**Figure 2A**) and a pair of mutagenesis primers, and could readily obtain the new mutant fragment #2 for all three viruses (**Figure 2B**). Successful rescue of the WA1 and Beta viruses, as evidenced by CPE, was consistently observed between day 3 and day 4, and the passage 0 (P0) stocks were typically harvested on day 4 or day 5 when CPE was >90%. The use of Vero E6-TMPRSS2 cells ensured the integrity of the furin cleavage site, as confirmed by sequencing of independently-rescued viruses (**Figure 2C**). The CPER-derived recombinant viruses also displayed the same plaque morphology as their parental isolates (**Figure 2D**), and the P0 virus titers consistently reached ~10^6^ PFU/mL (**Figure 2E**).

**FIGURE 2.**
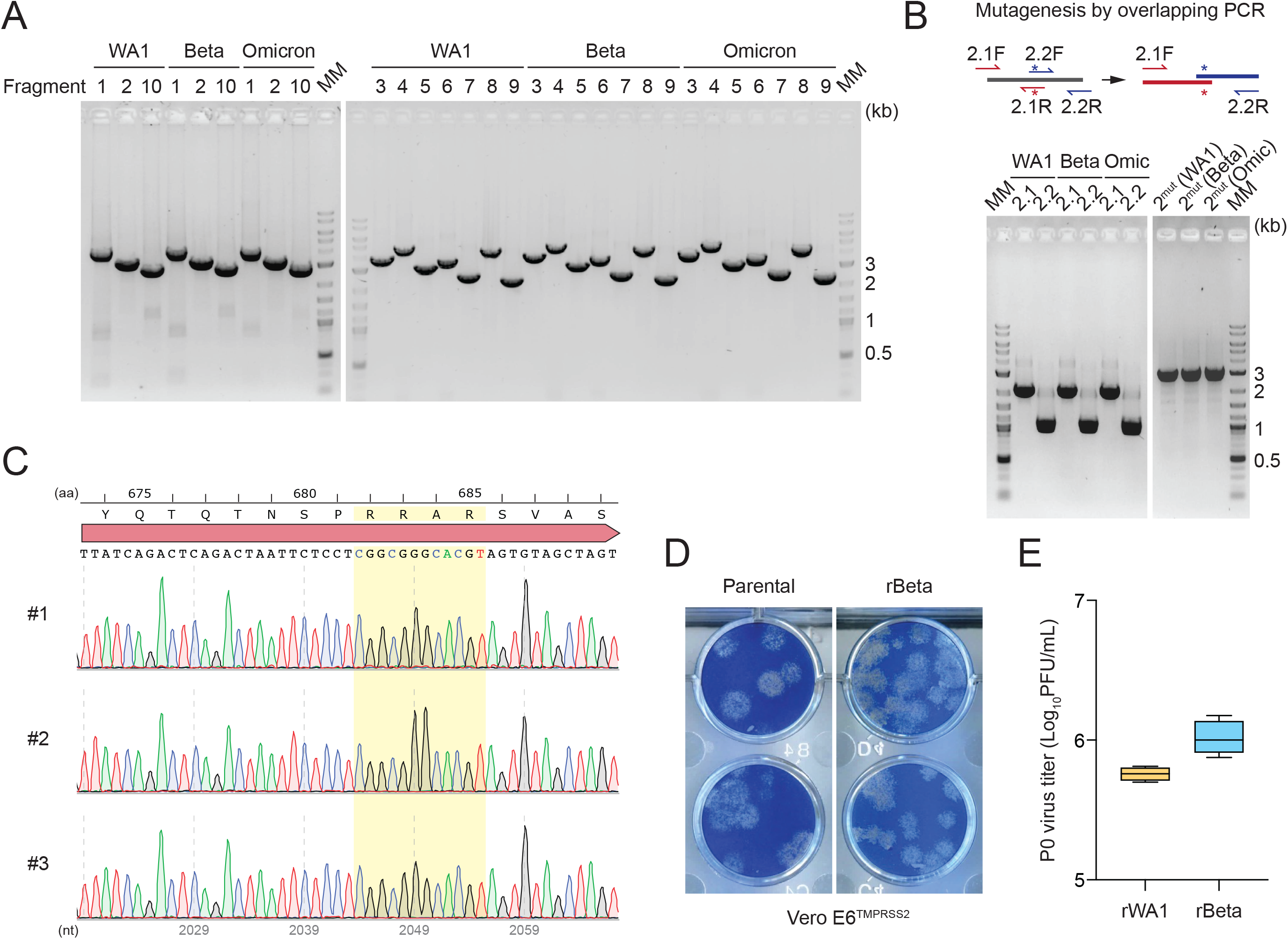
Cloning-free generation and characterization of CPER-derived recombinant SARS-CoV-2. (**A**) Representative gel images of the overlapping cDNA fragments amplified from purified SARS-CoV-2 genomic RNAs of the indicated virus strains. The primer sets described by Torii *et al*. [18] conform to the genome sequences of the ancestral strain (WA1) and the selected Beta and Omicron variants of concern (VOCs) with 100% complementarity. MM, molecular marker. (**B**) Schematic of the overlapping PCR strategy for site-directed mutagenesis in fragment 2 by using purified PCR product as a template (top panel), as well as representative gel images of the intermediate (2.1 and 2.2) and final (2^mut^) PCR products (bottom panel). MM, molecular marker. (**C**) Sequencing confirmation of the integrity of the spike furin cleavage site of the passage 0 (P0) virus stocks from three independent virus rescues using the optimized CPER approach. aa, amino acids; nt, nucleotides. (**D**) Plaque morphology on Vero E6-TMPRSS2 cells of recombinant Beta (rBeta) generated by optimized CPER as well as of its parental virus. (**E**) Virus titers of the P0 stocks of CPER-derived recombinant WA1 (rWA1) and rBeta, collected at day 5 and day 4 post-transfection of the CPER product, respectively *(n* = 4).

## DISCUSSION

CPER-based approaches offer considerable advantages over other reverse genetics systems for engineering positive-strand recombinant viruses harboring large genomes of >10 kb. First, they are PCR-based and better preserve viral genome sequences than plasmids or large DNA constructs (*i.e*. BACs) which require bacterial amplification. Second, CPER allows for manipulation of viral genome sequences via flexible PCR strategies with high accuracy, enabling rapid and reliable generation of recombinant mutant infectious clones for functional analysis of viral genes and specific motifs.

The herein-reported optimized CPER system, which was developed as part of our continuous efforts to define the role of SARS-CoV-2 genes in innate immune evasion (in particular, Nsp3 and its PLpro de-ISGylation activity) ([26] and Gack lab, unpublished data), addressed key limitations of the traditional CPER approaches that can compromise the robustness and efficiency of SARS-CoV-2 rescue. We provided proof of concept that, with the implementation of additional or modified steps – 5’ end phosphorylation, nick sealing, direct transfection into permissive cells – and through the use of a modified linker plasmid, SARS-CoV-2 rescue can be accelerated. At this point, we have not yet systematically determined which one(s) of these specific steps is functionally most important for the CPER optimization. It is conceivable that the combination of the new practices leads to successful virus rescue in a short time.

This optimized approach allowed for the accurate generation of reporter viruses and recombinant VOC strains, which displayed similar replication capacities as their respective parental viruses. The described optimization steps may be readily adapted also to other positive-strand RNA viruses such as other coronaviruses or alphaviruses, flaviviruses, and noroviruses. Further optimization of the reported workflow may be achieved by combining CPER and nick ligation in one reaction and by using other permissive cells (*e.g*. Caco-2) for transfection of the CPER product. Moreover, although the genome segmentation scheme and primer sets used in our studies (previously reported by Torri *et al.*) conform to the genome sequences of the selected Beta and Omicron strains, further optimization of the fragment scheme and primer locations could be attempted, considering phylogenetic analysis of sequence conservation, to achieve a universal set of primers that can be applied to all VOCs and emerging viral strains. It is also important to deep-sequence CPER-derived recombinant viruses and those generated by other reverse genetics systems, which would allow comparing the overall fidelity of different virus rescue approaches.

Our optimized CPER method may promote the functional analysis of recombinant viruses to evaluate viral determinants of pathogenesis, immune evasion and transmission. It could also be useful for the efficient generation of replication-’crippled’ viruses that may serve as live-attenuated vaccines with potentially higher efficacy than currently available COVID-19 vaccines. The optimized CPER approach described herein may also facilitate the incorporation of mechanism-based mutations that serve as built-in safety features (*e.g*. mutations in the Nsp1 gene and transcriptional regulatory sequence (TRS) [28, 29]) when studying certain viral variants or mutants.

### Additional comments regarding ethics and biosafety

Safe handling of viral agents such as SARS-CoV-2 is of utmost importance. Work with SARS-CoV-2 including recombinant viruses engineered using CPER (or other reverse genetics) approaches requires adequate biosafety biocontainment and is subject to institutional, local and/or federal regulations. Considering the ongoing debates about the dissemination of methods for reverse engineering of SARS-CoV-2 (see for example [30]), we consciously described in detail only the newly developed optimization steps of the CPER method, while mostly referring to published reports for the other steps of the CPER approach.

## MATERIALS AND METHODS

### Biosafety

SARS-CoV-2 genomic RNA extraction, cDNA synthesis, CPER transfection, and live virus experiments were all conducted in the BSL-3 facility of the Cleveland Clinic Florida Research and Innovation Center (CC-FRIC). Sterility-tested viral cDNA was handled in a BSL-2 laboratory following standard biosafety practices and procedures. All work was reviewed and approved by the CC-FRIC Institutional Biosafety Committee in accordance with the National Institutes of Health (NIH) Guidelines.

### Cells and viruses

Vero E6 (#CRL-1586) and HEK293T (#CRL-3216) cells were purchased from the American Type Culture Collection (ATCC) and were maintained in Dulbecco’s modified Eagle’s medium (DMEM, Gibco) supplemented with 10% fetal bovine serum (FBS, Gibco), 2 mM L-Glutamine (Gibco), 1 mM sodium pyruvate (Gibco) and 100 U/mL of penicillin–streptomycin (Gibco). Vero E6 cells stably expressing human TMPRSS2 were generated by lentiviral transduction followed by selection with blasticidin (40 μg/mL; Invivogen). SARS-CoV-2 strains hCoV-19/USA-WA1/2020 (NR-52281), hCoV-19/USA/MD-HP01542/2021 (Lineage B.1.351; Beta variant) (NR-55282), and hCoV-19/USA/MD-HP20874/2021 (Lineage B.1.1.529; Omicron variant) (NR-56461) were obtained from BEI Resources, National Institute of Allergy and Infectious Diseases (NIAID), NIH.

### Viral genomic RNA purification and first-strand cDNA synthesis

Viral genomic RNA was purified from 280 μL virus-containing media using the QIAamp Viral RNA Mini Kit (Qiagen) as per the manufacturer’s instructions and eluted in 60 μL nuclease-free water. Reverse transcription for first-strand cDNA synthesis was performed by using the LunaScript RT SuperMix Kit (NEB) containing both oligo(dT) and random primers in a reaction consisting of 10 μL genomic RNA, 4 μL 5× SuperMix and 6 μL nuclease-free water with the cycling condition as follows: 2 min at 25°C, 20 min at 55°C, and 1 min at 95°C. One microliter of RNase H (5 U; Thermo Scientific) was subsequently added and the reaction mix was incubated at 37°C for 20 min.

### DNA constructs

The bacterial artificial chromosome (BAC) encoding a GFP reporter SARS-CoV-2 in the background of hCoV-19/Germany/BY-pBSCoV2-K49/2020 (GISAID EPI_ISL_2732373) was kindly provided by Armin Ensser (Friedrich-Alexander University Erlangen-Nürnberg, Germany) and has been described previously [11]. pGL-CPERlinker was assembled from synthetic DNA oligonucleotides and fragments (IDT) as well as the ampicillin resistance cassette and the origin of replication derived from pUC19 (NEB).

### CPER reaction and transfection

To amplify the 10 viral cDNA fragments (either from BAC or the first-strand viral genomic cDNA), previously reported primer sets were used [18]. The primers for amplification of the linker fragment from pGL-CPERlinker are: GL-CPERlinkF (5’-CTTAGGAGAATGACAAAAAAAAAAAAAAAAAAAAAAAAAAAGGCCGGCATGGTCCCAGCC-3’) and GL-CPERlinkR (5’-GTTACCTGGGAAGGTATAAACCTTTAATACGGTTCACTAAACGAGCTCTGCTTATATAG-3’). Amplification of each fragment was carried out by using the PrimeSTAR Max DNA polymerase (Takara Bio) in a 50 μL PCR reaction containing 0.2 μM each primer and 1 ng BAC or 2 μL viral cDNA as the template with the cycling condition as follows: 10 s at 98°C; 35 cycles of 10 s at 98°C, 5 s at 55°C, 25 s at 72°C; and 2 min at 72°C. All PCR products were gel purified by using the Monarch DNA Gel Extraction Kit (NEB) and eluted in 20 μL nuclease-free water. The purified fragments were then 5’ phosphorylated in a 50 μL reaction containing 10 U of T4 polynucleotide kinase (NEB) and cleaned up through the Monarch PCR & DNA Cleanup spin columns (NEB). CPER assembly was performed as previously described by combining 0.05 pmol of each fragment in a 50 μL reaction containing 2.5 U PrimeSTAR GXL DNA polymerase (Takara Bio) and using the ‘condition 3’ cycling parameters [18]. Immediately before transfection, the CPER product was subject to post-PCR nick sealing for 30 min at 50°C and 30 min at 60°C in a 25 μL reaction containing 1 mM β-nicotinamide adenine dinucleotide (NAD+) (NEB) and 0.5 μL HiFi Taq DNA ligase (NEB). The final CPER product was transfected into Vero E6-TMPRSS2 cells seeded into 6-well plates (~ 5 × 10^5^ cells per well) by using the *Trans*IT-X2 Dynamic Delivery System (Mirus Bio) as per the manufacturer’s instructions. After 24 hours, the culture media was replaced with DMEM containing 2% FBS, 2 mM L-Glutamine, 1 mM sodium pyruvate, 1× non-essential amino acids (Gibco), 10 mM HEPES (Gibco), and 100 U/mL of penicillin–streptomycin. For the classical CPER method, the unsealed CPER product was first transfected into HEK293T cells by using *Trans*IT-LT1 (Mirus Bio), and the trypsinized cells were then overlaid onto Vero E6-TMPRSS2 cells at 6 hours post-transfection, as previously described [17].

### Virus titration and sequencing

The titers of the P0 virus stocks were determined by plaque assay. Briefly, a monolayer culture system of Vero E6-TMPRSS2 cells was incubated with ten-fold serially diluted virus-containing media. The inoculum was removed after 2 hours, and the cell monolayers were washed twice with PBS and then overlaid with 1% colloidal microcrystalline cellulose (Sigma) in MEM containing 2% FBS, 2 mM L-Glutamine, 1× non-essential amino acids, 10 mM HEPES, and 100 U/mL of penicillin–streptomycin. Plaques were visualized by Coomassie Blue staining on day 3. For P0 virus sequencing, viral genomic RNA was purified and the first-strand cDNA was synthesized as described above. Nine fragments encompassing the whole genome [31] were then amplified from the cDNA and subsequently subjected to Sanger (Azenta Life Sciences) or Nanopore sequencing (Plasmidsaurus).

## AUTHOR CONTRIBUTIONS

G.L. designed and performed all experiments and analyzed the data. M.U.G. supervised the study. G.L. and M.U.G. wrote the manuscript.

## ACKNOWLEDGEMENTS

This study was funded, in part, by the US National Institutes of Health grant R37 AI087846 (to M.U.G.). We greatly thank Armin Ensser (Friedrich-Alexander University Erlangen-Nürnberg) for providing reagents.

